# Species invasiveness and community invasibility of US freshwater fish fauna revealed via trait-based analysis

**DOI:** 10.1101/2022.03.04.481515

**Authors:** Guohuan Su, Adam Mertel, Sébastien Brosse, Justin M. Calabrese

**Affiliations:** Center for Advanced Systems Understanding (CASUS), Görlitz, Germany; Helmholtz-Zentrum Dresden-Rossendorf (HZDR), Dresden, Germany; Laboratoire Evolution et Diversité Biologique (EDB), Université de Toulouse, CNRS, IRD, UPS, Toulouse, France; Department of Biology, University of Maryland, College Park, MD, USA

## Abstract

While biological invasions are recognized as a major threat to global biodiversity, determining species’ abilities to invade new areas (species invasiveness) and the vulnerability of those areas to invasions (community invasibility) are still poorly understood. Here, we used trait-based analysis to profile invasive species and quantify the community invasibility for >1,800 North American freshwater fish communities. We show that species with higher reproduction rates, longer life spans and larger sizes tend to be more invasive. Community invasibility peaked when the functional distance among native species was high, leaving unoccupied functional space for the establishment of potential invaders. Invasion success is therefore governed by both the functional traits of non-native species determining their invasiveness, and by the functional characteristics of the invaded community determining its invasibility. Considering those two determinants together will allow better predictions of invasions.

## Introduction

Freshwater systems are among the most threatened ecosystems and most of the world’s river basins have been severely altered by human activities (*1*, *2*). Among them, habitat fragmentation and non-native fish introductions are the most pervasive (*2*). In particular, fish introductions have markedly changed fish community structure and composition in rivers worldwide (*3*, *4*). Nevertheless, our ability to predict invasions remains meager considering both non–native species’ invasiveness (i.e., the capacity of a species to colonize areas where it does not naturally belong), and native communities’ invasibility (i.e., the vulnerability of native communities to non-native species establishment) (*5*) are poorly understood properties.

Invasiveness has frequently been assessed by comparing functional traits or life history strategies between non-native and native species from the recipient communities (e.g., *6*, *7–10*). Non-natives have been reported to belong to higher trophic levels and have distinct swimming capacities compared to natives (*10*, *11*). However, few studies have tested which among these differential traits actually help a species to colonize areas where it does not naturally belong. In other words, while it is clear that invasive species often feature different traits, the relationship between functional traits and species invasiveness *per se* remains to be explored.

Moreover, species invasiveness is likely coupled with the functional traits of recipient communities (*12*), which could determine their vulnerability to non-native species establishment. Two mutually exclusive ecological hypotheses have been frequently invoked to explain community invasibility. First, the biotic acceptance hypothesis predicts that the number of successful invasive species is positively related to native species richness in the recipient community, as favorable environmental conditions sustaining high native species richness should also benefit non-native species (*13*). In contrast, the biotic resistance hypothesis predicts a negative relationship between native and non-native species richness, because competitive interactions between native and non-native species will increase with native species richness, thus excluding most non-native species (*14*). However, neither of these two hypotheses explained non-native species richness in river basins around the world, and only anthropogenic disturbances – non-native species releases and environmental degradation – were responsible for increased non-native species richness (*4*).

The lack of clear relationships between recipient community properties and invasibility might stem from the use of taxonomic diversity metrics such as richness or identity of the species. Those metrics may not accurately predict invasibility because the diversity in species does not predict the diversity of the functions they support (*15*). In fish communities most species are functionally redundant, whereas a few have unique functional traits (*16*, *17*). Such functional uniqueness makes the communities and the functions they support vulnerable to environmental changes, implying that so-called ecosystem insurance (*18*) is only true for a few redundant functions. Thus, community invasibility might not be explained by the diversity of the functions experienced by a community but by the functional redundancy among species from the focal community. Communities with functions supported by unique species, should therefore be more vulnerable to invasions than communities with strong functional packing.

Our aim was therefore to characterize the functional structure of communities by considering the range (e.g., functional richness) and the partitioning (e.g., functional evenness, functional divergence) of functions within each community (*15*, *16*). Understanding whether the functional structure of local communities and functional similarity (or distinctness) between non-native and native species affect the invasion process could help identify whether community invasibility is primarily governed biotic acceptance or by biotic resistance. For instance, the unsaturation of local communities ensures establishment of new species without native species exclusion (*19*, *20*), thus suggesting a dominance of biotic acceptance. But even in unsaturated communities, if the introduced non-native species are functionally similar to natives it might still generate competitive effects, and lead to biotic resistance (*16*, *21*). We therefore predict that the functional similarity between non-native and native assemblages, as well as the functional structure of local communities constitute important determinants of the invasion process.

Here, we examined how functional trait analysis can unify the species-centered and community-focused views of the invasion process and yield new insights into both the invasiveness of particular species, and the invasibility of recipient communities. We used fish occurrence data from more than 1,800 watersheds across the United States coupled with 20 fish life-history traits (i.e., morphological, physiological, behavioral) to compute two distance metrics between nonnative and native species (Fig. 1), and six complementary functional diversity indices for recipient fish communities at the watershed level. Our goal is not only to profile the functional characteristics of invaders, but also to quantify the vulnerability of recipient communities based on their functional attributes. We therefore expect that the invasion risk posed by a non-native species results from the combination of its own functional attributes and of the functional characteristics of the recipient community.

**Fig.1.**
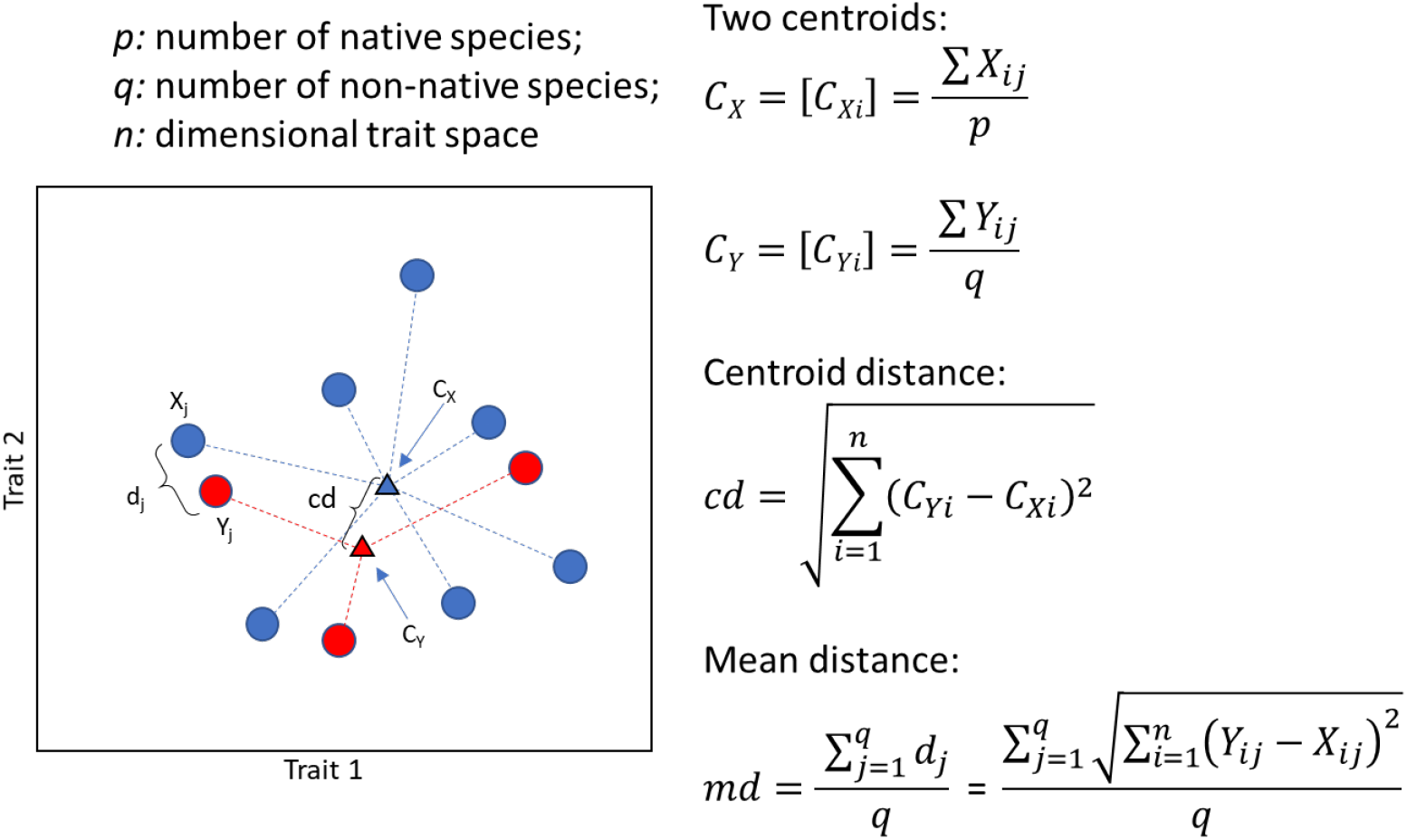
An example showing how centroid distance (*cd*) and mean distance (*md*) are computed. The ***p*** and ***q*** individual native and non-native species in a ***n***-dimensional trait space (here ***n=2***) are represented by blue and red circles. Vectors ***Y_j_*** and ***X_j_*** represent the positions of non-native species ***j*** and its nearest native species ***j***. ***d_j_*** is the distance between ***X_j_*** and ***Y_j_***. ***C_X_*** and ***C_Y_*** (triangles) are the centroids of the ***p*** native species and ***q*** non-native species. In that case, ***C_X_* = [C_Xi_]** and ***C_Y_* = [C_Yi_]**, where ***C_Xi_*** and ***C_Yi_*** are the mean value of trait ***i*** for native and non-native species ***cd*** is the distance between ***C_X_*** and ***C_Y_***. ***md*** is the mean distance between all non-native species and their nearest native neighbors.

## Results

### Spatial distribution of non-native species

Among the 1,873 considered watersheds, covering most of the continental US, we identified 1,720 watersheds that have at least one record of a non-native species. 1,560 watersheds that had translocated species (i.e., species that were native to the continental US but translocated to watersheds from which they were historically absent), and 1,353 watersheds that had exotic species (i.e., species that were historically absent from the continental US). The 1,873 focal watersheds contain 862 fish species, including 562 native species that have never invaded other watersheds, 227 translocated species, and 73 exotic species. Native and non-native species richness spatial patterns were contrasted, with the highest native fish richness in the watersheds belonging to the Mississippi drainage, whereas non-native species richness peaked east and west of the Mississippi (Fig. 2).

**Fig.2.**
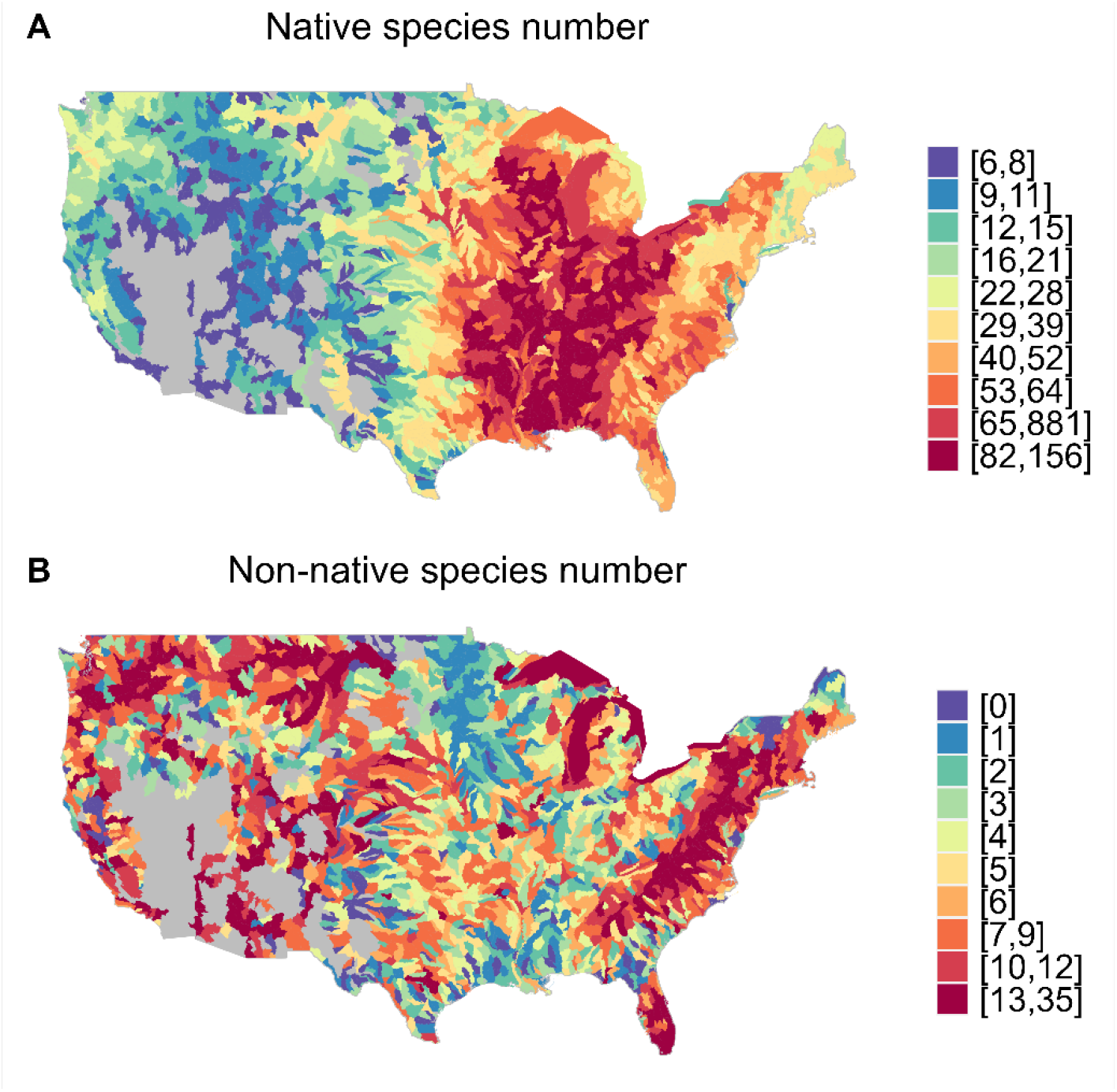
Number of native and non-native species in the 1,873 watersheds in the US. A) native species; B) non-native species.

### Functional difference between native and non-native fishes

Both translocated and exotic species showed different distributions along most of the 20 functional traits from native species. Moreover, some of the traits’ average values changed gradually from native, to translocated and to exotic species (Fig. S1, S2). Indeed, among the 10 morphological traits, values of maximum body length and relative eye size gradually increased, while body elongation gradually decreased from native, to translocated, and to exotic species. Among the other 10 ecological and life-historical traits, average values of longevity, percent of euryhaline species, and percent of diet breadth also showed gradually increasing trends across the three groups. The translocated and exotic species both showed higher fecundity than the native species (K-S test, *P* < 0.001). However, parental care for exotic species is significantly more frequent than for the native and translocated species (Chi-square test, *P* < 0.001)., while the latter two groups did not differ (Fig. S1, S2).

### Species invasiveness

Species invasiveness was significantly influenced by a set of five predictors, each contributing more than 6.25% in the boosted regression trees model (hereafter BRT). According to the cross-validation procedure, the model explained 34.7% of the total deviance. Partial dependency plots in Fig. 3 showed that the PCA axis (Repro_PCA1) combining maximum body length, fecundity, longevity and age at maturity contributed the most (39.2%) to invasion frequency, followed by body elongation (17.7%), diet breadth (10.5%), trophic level (9.8%) and the PCA axis (Temp_PCA1, 7.4%) combining the three temperature variables. Neither the other traits nor the type of the non-native species had a significant influence on the patterns (Fig. 3). Despite a few outliers, invasiveness is positively correlated to the reproduction-related traits, diet breadth, and trophic level. However, invasiveness reached its highest values at mid-high trophic levels and then decreased slightly thereafter. In contrast, the invasiveness is negatively correlated with body elongation and temperature-related traits (Fig. 3).

**Fig.3.**
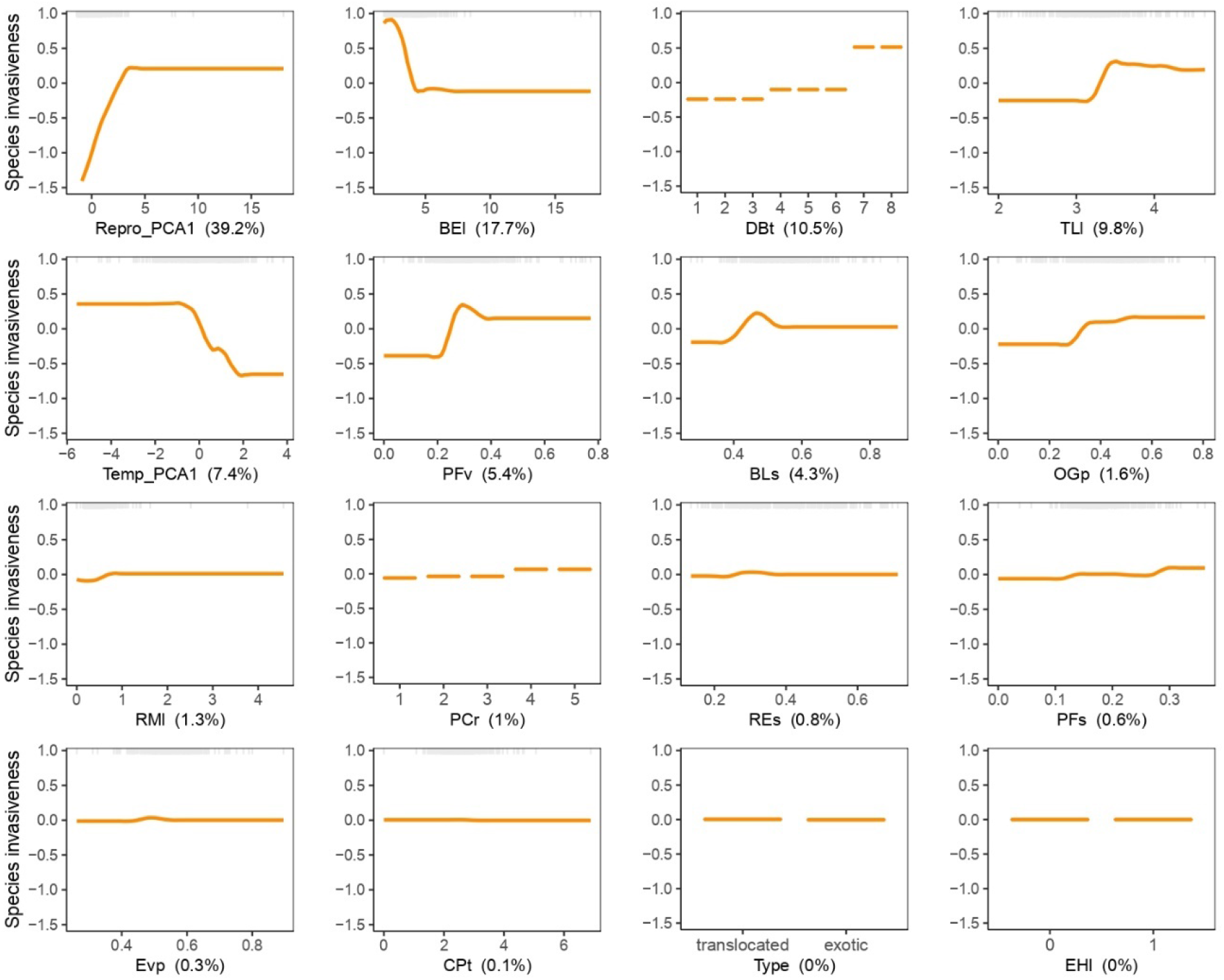
Results of boosted regression trees showing the partial dependency between species invasiveness and fish functional traits. Repro_PCA1: first PCA axis of maximum body length, longevity, fecundity, and mature age traits; BEl: body elongation; DBt: diet breadth; TLl: trophic level; Temp_PCA1: first PCA axis of temperature range, minimum and maximum traits; PFv: pectoral fin vertical position; BLs: body lateral shape; OGp: oral gape position; RMl: relative maxillary length; PCr: parental care; REs: relative eye size; PFs: pectoral fin size; EVp: eye vertical position; CPt: caudal peduncle throttling; EHl: euryhaline. The value in parentheses in each panel shows the percentage of contribution of each trait considered in the model, and contributions > 6.26% are significant.

### Community invasibility

The BRT model explained 67.6% of the total deviance for the patterns of community invasibility. Partial dependency plots presented in Fig. 4 show the effect of a particular variable on the invasibility after accounting for the average effects of all other variables in the model. Fitted functions by the BRT model were frequently nonlinear and varied in shape. Six variables related to trait-based distance, functional structure, environment and human activities were found to be the best predictors of community invasibility, with relative contributions ranging from 9% to 17% (Fig. 4). Among them, the functional distinctness between non-native species and native fish assemblages, measured as the centroid distance between the non-native and native species assemblages, was the most influential predictor (16.8%) and was negatively related to community invasibility. In contrast, mean functional distance between non-native species and their nearest neighbors (9.6%) was positively related to the community invasibility. Functional specialization (9.8%) was found to be the most influential among the six functional diversity indices and had a positive influence on community invasibility. As expected, number of invasions increased significantly with the intensity of human activities, especially for the human footprint variable (9.1%, Fig. 4).

**Fig.4.**
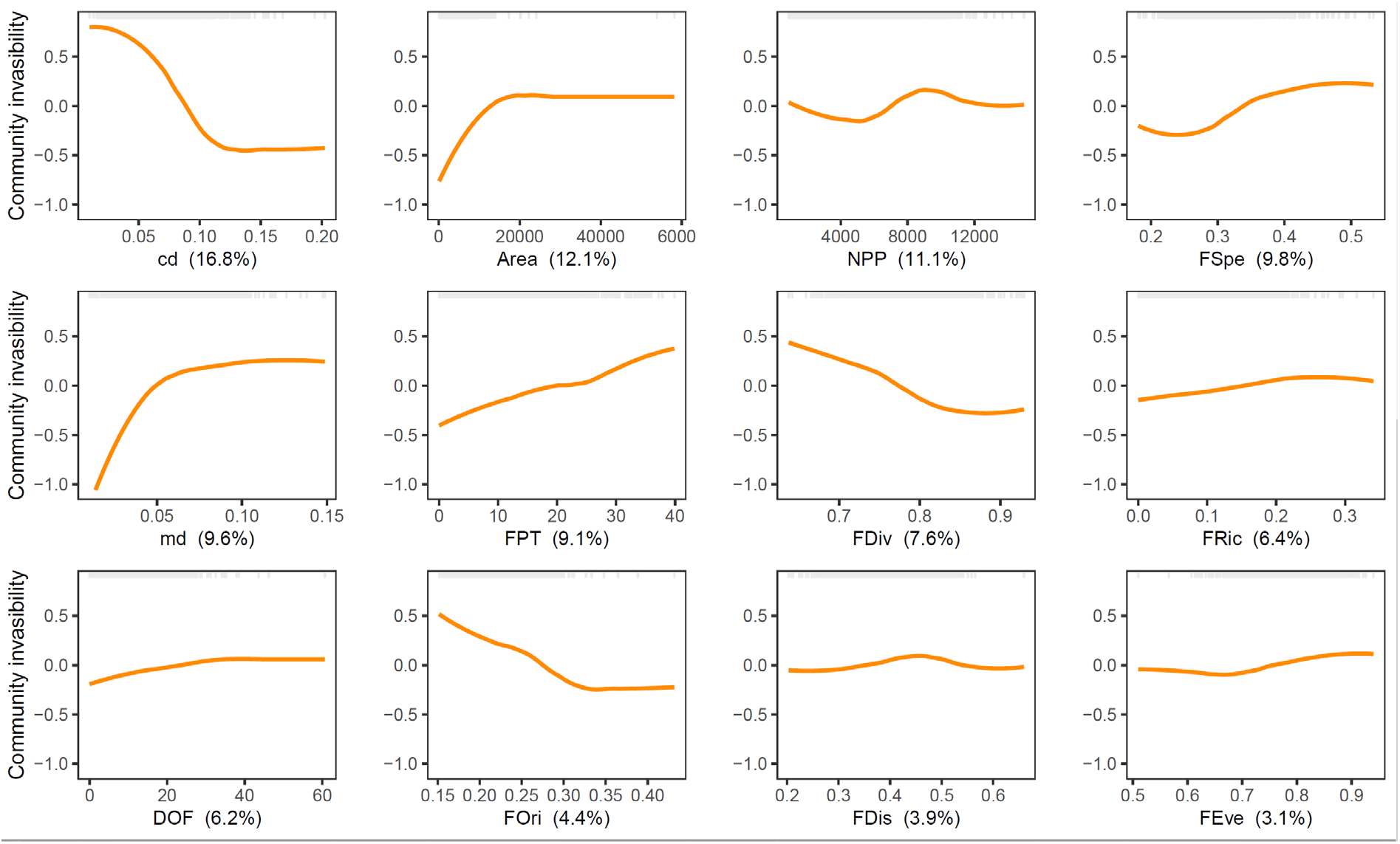
Results of boosted regression trees showing the partial dependency between community invasibility and predictors related to recipient community functional structure, distance between invasive and native species, environment, and human activities. cd: centroid distance; Area: watershed area; NPP: net primary productivity; FSpe: functional specialization; md: mean distance; FPT: human footprint; FDiv: functional divergence; FRic: functional richness; DOF: degree of river fragmentation; FOri: functional originality; FDis: functional dispersion; FEve: functional evenness. The value in parentheses in each panel shows the percentage of contribution of each predictor considered in the model, and contributions > 8.33% are significant.

## Discussion

Fish morphological traits are different between native and non-native species. Specifically, studies considering morphological differences between natives and non-natives at the river basin scale showed that non-native species had larger size and less elongated bodies than their native counterparts. This general trend holds throughout the globe, and indicates that most non-native species are adapted to live in lentic or slow flowing habitats (*8*, *17*, *22*). Within river basins, (*9*) also revealed that established non-native fish species exhibited distinct life-history strategies compared to the native species in the Colorado River basin. Our regional (watershed) approach over the continental US revealed that functional traits, including morphological, physiological and life-historical aspects differ between native and non-native species (including translocated and exotic species), therefore paralleling previous studies at both local and global scales (*2*, *9*, *10*).

However, our findings also show that not all of the traits differing between native and non-native species contribute to the invasiveness of non-native species. Instead, species’ invasiveness was only predicted by a few traits, among which high reproduction rates, long life spans and large sizes are the most influential. High reproduction rate is considered as an advantage to establish and spread in a novel environment, as shown by (*23*) for the common carp (*Cyprinus carpio*) invasion in Australia. Moreover, life span and body size are positively linked to the dispersal ability of the species, and therefore favor the post establishment spread of large and long-lived species (*7*, *22*). In addition, large species are often preferred for aquaculture and angling, which are among the most efficient pathways of introduction, and generate massive and widespread fish releases in natural environments (*24*, *25*) that contribute to the invasiveness of those species.

In contrast, although other traits of non-native species also significantly differed from those of native species, their contribution to species invasiveness remained negligible. This could be explained by the complexity of the invasive process (including introduction, establishment and spread steps, each determined by distinct drivers) and the variety of factors that might affect invasion success (including difficult to measure factors such as propagule pressure or temporal dynamics of introduction) (*26*, *27*). Indeed, (*17*) revealed that environmental filtering acts together with human preference for particular fish morphologies to jointly determine the morphological characteristics of introduced non-native species. Intriguingly, our results show that top predators with a relatively narrow diet breadth, are not among the best invaders. Instead, omnivorous fish with a wide diet breadth appear to be more invasive. For instance, top predators feeding only on fishes such as peacock bass (*Cichla ocellaris*, exotic) and Muskellunge (*Esox masquinongy*, translocated), have been recorded to invade 4 and 25 watersheds in the US, respectively. In contrast, the common carp (*Cyprinus carpio*, exotic) and yellow bullhead (*Ameiurus natalis*, translocated), with a moderate trophic level but a wide diet breadth, have been recorded to invade 1,131 and 452 watersheds in the US, respectively. In addition, although the distributions of most traits significantly differed between the translocated and exotic species (table S2), the type of invasion (exotic/translocated) did not influence the species invasiveness models, demonstrating that species identity or native origin is a poor proxy of invasiveness compared to functional traits.

Community invasibility of US watersheds was more influenced by the functional structure of the communities than by human disturbances. Surprisingly, human footprint and the degree of river fragmentation had only small effects on community invasibility, whereas they are recognized as major predictors of the number of non-native species (*4*, *28*, *29*). For instance, (*2*) revealed that patterns of change in global fish biodiversity were dominated by the introduction of non-native species in anthropized areas with high human footprint and intense river fragmentation. Our regional findings contrast with these global results as we report a weak influence of human activities on community invasibility patterns across the US watersheds. This inconsistency might be rooted in the different spatial scales considered. At the watershed scale, the measurement of river fragmentation only accounts for dams located on the focal watershed, while hydrologically mediated effects might spread far downstream (*30*). Our findings also tend to support the biotic acceptance hypothesis, which is verified by the positive correlation between the watershed area (and thus the native species richness) and invasibility, therefore confirming that species rich native assemblages are not insured against non-native species establishment.

More importantly, by considering the functional diversity metrics of the local community and the trait-based distance metrics between local and non-native assemblages, we were able to highlight a novel mechanism explaining community invasibility. The highest invasibility was indeed recorded when native and non-native species pools from a watershed share the same functional diversity (low centroid distance between non-native and native species assemblages), and individual native and non-native species within that assemblage are not functionally redundant (high mean distance between the non-native species and their nearest neighbors). This indicates that most non-native species pack into the center of the native species’ functional space, but keep distance from their native neighbors, thereby avoiding competitive interactions that are prone to reduce the chances of establishment. Such a process can therefore be viewed as an environmental filtering effect that increases the overall functional similarity between native and non-native species pools in the same watershed (thus resulting in an apparent biotic acceptance effect). Then, among the species that successfully passed the environmental filter, only those occupying an available functional niche can establish and not suffer from biotic resistance. The interplay between environmental filtering at the community scale and biotic resistance at the species scale therefore represents a novel process that may solve the long-standing debate about the environmental vs biotic determinants of biological invasions (e.g., *4*, *31*). We here show that both processes act together but the former at the community level and the latter at the species level. Thus, communities with lower density in species in their functional center will be the most sensitive to invasions, which is confirmed by the positive correlation between community invasibility and functional specialization. Indeed, functional specialization represents the proportion of generalist species (i.e., species close to the center of the functional space, (*16*)) in a community, and high functional specialization indicates that more gaps are available around the functional center. In contrast, the size of the functional space (i.e., functional richness) and other metrics representing functional structure of assemblages do not facilitate invasions. At least for the US fish fauna, functional specialization can be considered as proxy for vulnerability to non-native species (i.e., community invasibility), and it could therefore be of major interest when designing management actions to avoid further invasions of the most sensitive watersheds. This is of particular importance given the current spread of non-native species throughout the world, as well as the predicted emergence of new invaders (*32*, *33*). We thus implore future studies to evaluate the relevance of the functional specialization metric as a proxy of invasion vulnerability in other regions and on other taxa.

To conclude, our study shows the importance of functional traits in the analysis of species invasiveness and community invasibility. We confirm that functional differences between native and non-native fish species exist, but species invasiveness is dominated by only a few functional traits among them. Our results also provide new insights into the mechanisms promoting community invasibility. Though our findings tend to support the human activity and biotic acceptance hypotheses, in essence, community invasibility cannot be simply driven by one of them or by their joint effect. Instead, the original mechanisms unveiled in our study suggest that functional similarity between the non-native and native species and the local community functional structure are more influential than human activity and biotic acceptance in shaping community invasibility patterns. Communities with higher levels of functional redundancy or denser functional centers would have stronger resistance to invasive species (i.e., lower invasibility). This could explain why the speciose fish communities in the Amazon river basin, which are highly redundant in functions (*34*, *35*), have received few non-native species (*36*). Our study also raises a new question as to why most species, whether native or non-native, tend to gather in or invade the crowded center of the functional space, even if the space near the border is almost empty. We expect that the distribution of resources and the relative position of species niches are the key factors, but further evidence is needed to confirm this.

## Materials and Methods

### Species occurrence data

Native species occurrence records (804 species considered) at the watershed scale (i.e., Hydrologic Unit Code 8; HUC8) were obtained through NatureServe (https://www.natureserve.org) and included both extant and extinct species to account for species historically present in a given watershed but extirpated as a potential consequence of various human activities such as species invasions. Occurrence records of the naturalized or established non-native fish species (321 species considered) were obtained through the U.S. Geological Survey (USGS) Non-indigenous Aquatic Species (NAS) database (http://nas.er.usgs.gov) at the watershed scale (i.e., Hydrologic Unit Code 8; HUC8). Non-native species data includes exotic species that were historically absent from the continental US, and translocated species that were native to the continental US but translocated to watersheds from which they were historically absent. This dataset only considered the identified species that locally create self-sustaining populations, thus we excluded records of non-self-sustaining or eradicated populations, vagrant species detected in only one sampling occasion, and non-identified or hybrid species.

### Functional traits

We collected ten morphological traits related to fish locomotion and food acquisition and ten additional traits related to life-history and physiological functions from FISHMORPH database (*37*), the Fish Traits database for North American freshwater fishes (*38*), and FishBase (*39*). The ten morphological traits are maximum body length, body elongation, relative eye size, oral gape position, relative maxillary length, vertical eye position, body lateral shape, pectoral fin vertical position, pectoral fin size, and caudal peduncle throttling. The ten additional ecological traits are longevity, fecundity, mature age (i.e., age of sexual maturity), trophic level, temperature range, minimum temperature, maximum temperature, euryhaline (yes, no), parental care (non-guarders1: open substratum spawners, non-guarders2: brood hiders, guarders1: substratum choosers, guarders2: nest spawners, bearers), and diet breadth (from 1 to 9). See Table S1 for details on the 20 functional traits.

Due to insufficient information on some species, some values were missing in the raw functional trait data. Overall, 21% of the values were missing in the raw trait dataset of 959 fish species. We statistically imputed these missing values (NA) with a machine learning algorithm called ‘missForest’ (*40*, *41*). This method uses a random forest trained on the observed values of a data matrix to predict the missing values and automatically calibrates the filling values by a set of iterations. In the imputation process, after each iteration the difference between the previous and the new imputed data matrix is assessed for the continuous and categorical parts, and the algorithm stops once both differences become larger (*40*). It can be used to impute continuous and/or categorical data and is not biased by complex interactions or nonlinear relationships. We included the evolutionary relationships between species in the imputation process by including the first ten phylogenetic eigenvectors in the matrix to be imputed (*42*). We tested the accuracy of this method in filling in missing values on a random set of 350 species with complete values for all traits. We randomly deleted 20% of the values for the 350 species, and then imputed them with missForest. We then compared the simulated values to the actual values, and repeated this procedure 100 times. Finally, we quantified imputation accuracy by calculating the Spearman correlation coefficient between the actual and imputed data, which varied from 0.86 to 0.95. In contrast, the classical imputation method of filling in missing observations with the average trait value of the 80% species with data, produced average correlation coefficients that ranged from 0.79 to 0.85, confirming the improved performance of missForest (Fig. S3).

### Predictors used in the invasibility models

#### a. Functional diversity indices

First, we calculated trait dissimilarity between species pairs in the communities using the Gower pairwise distance (*43*). This metric can handle multiple types of data (e.g., categorical, ordinal and continuous traits). We then used principal coordinate analysis (PCoA) to build the functional space on the first 5 principal coordinate axes, which explained over 80% of the total variance. We removed watersheds with fewer than 6 species to meet the criteria for calculating functional diversity indices, which resulted in 1,873 watersheds for the following analyses. Then we computed six complementary functional indices that are frequently used in functional diversity studies (*16*, *44–46*): functional richness (FRic), functional evenness (FEve), functional divergence (FDiv), functional dispersion (FDis), functional specialization (FSpe), and functional originality (FOri). These six metrics were used to represent the functional size and structure of the recipient fish community in each watershed.

FRic measures the size (i.e. convex hull) of the functional space; FEve measures the regularity of traits in the functional space; FDiv measures the proportion of species with the most extreme trait values; FDis measures deviation of species trait values from the center of the functional space; FSpe measures the mean distance of a species from the rest of the species pool; and FOri measures the distance between each species and its nearest neighbor (*16*, *47*).

#### b. Trait-based distance between invasive and native species

We computed two novel metrics to represent the distance between the non-native species assemblage and recipient community for each watershed in the five-dimensional functional space, which thus reflect the degree of functional redundancy between them. First, the centroid distance (*cd*) is the distance between the centroids of non-native and native species assemblages, which reflects the overall relative positions of the two groups. Second, the mean distance (*md*)is the distance between all non-native species and their nearest native neighbors, which reflects the average position of individual non-native species relative to their nearest neighbors. See Fig. 1 for details about how these two metrics were calculated.

#### c. Environmental and human related variables

We also included the variables widely used in testing the three main hypotheses relevant to community invasibility (*4*). Variables related to biotic acceptance/resistance hypotheses were selected as the native species richness (NSR), net primary productivity (NPP) and watershed area (Area). Variables related to human activity hypothesis were selected as human footprint (FPT), gross domestic product (GDP), and degree of fragmentation (DOF).

If community invasibility is strongly and positively correlated to NSR, NPP and Area, the biotic acceptance hypothesis will be supported. Otherwise, if community invasibility is strongly and negatively correlated to NSR, the biotic resistance hypothesis will be supported (*4*). Human activity hypothesis will be supported if community invasibility is highly correlated to the FPT, GDP and DOF.

NPP was taken from an online data repository (http://files.ntsg.umt.edu/), using the mean annual NPP from 2000 to 2015. FPT is a comprehensive representation of anthropogenic threats to biodiversity, which cumulatively accounts for eight human pressures—built environments, croplands, pasture lands, human population density, night lights, railways, major roadways, and navigable waterways. The FPT dataset (resolution: 1 km^2^) was taken from (*48*). GDP measures the size of the economy and is defined as the market value of all final goods and services produced within a region in a given period. The GDP dataset (1 square degree resolution) was taken from (*49*). DOF measures the degree to which river networks are fragmented longitudinally by infrastructure, such as hydropower and irrigation dams (*1*). The DOF dataset (resolution: 500 m2) was taken from (*1*).

We mapped NPP, FPT, GDP and DOF by their relative resolution grid data over the watershed-scale map and then calculated the mean value of all the cells covered by each watershed using QGIS.

### Statistical analysis

We compared the distributions of 20 traits among the three assemblages (i.e. native, translocated and exotic species) via the Kolmogorov–Smirnov test (hereafter K–S test) for continuous traits and the Chi-square test for categorical traits.

Since maximum body length, fecundity, longevity, and age at maturity are highly correlated (Pearson r > 0.7, Fig. S4A), we used principal component analysis (PCA) and chose the first PC axis as a combined reproductive trait (Repro_PCA1), which represents 71.9% of the total variance (Fig. S4B). Similarly, we did a PCA for the three temperature related traits and chose the first PC axis (Temp_PCA1), which represents 74.9% of the total variance (Fig. S4C). We used the invasion frequency (i.e., the frequency of occurrence of a species in the watersheds where it is not historically present) of each established non-native species across the 1,873 watersheds as a proxy of species invasiveness. Then, we employed boosted regression trees (BRT) to identify which functional traits or trait combinations determine species invasiveness. We also included the type of invasion (i.e., exotic or translocated) in the BRT model to test whether the different categories of non-native species behaved differently. Therefore, 16 predictors were considered in this BRT model. We applied the methodology proposed by (*50*) using a BRT model that assumes a Poisson distribution of the response variable.

We then quantified the correlations among the above predictors for the community invasibility model and found that NSR and GDP were highly correlated (Spearman rho>0.7, Fig. S5) with FRic and FPT, thus we removed NSR and GDP from the following models. We computed the established non-native species number for each of the 1,873 watersheds as the community invasibility, and applied the Poisson BRT model to assess the relative importance of each of the 12 above-described predictors on the observed invasibility of the watershed-level communities.

The BRT models were fitted using the ‘gbm.step’ function in ‘dismo’ package in R (*50*), which allows for the specification of four main parameters: bag fraction (*bf*), learning rate (*lr*),tree complexity (*tc*) and the number of trees (*nt*). *bf* is the proportion of samples used at each step, *lr* is the contribution of each fitted tree to the final model, *tc* is the number of nodes of each fitted tree determining the extent to which statistical interactions were fitted, and *nt* represents the number of trees corresponding to the number of boosting iterations. The optimal setting of the parameters was chosen using 10-fold cross validation (CV). The procedure provides a parsimonious estimate, CV – D^2^ (i.e., the cross validated proportion of the deviance explained), representing the expected performance of the model when fitted to new data (*50*). Using CV, we explored different combinations of the parameters to be set and retained the optimal model showing the highest CV – D^2^. We used the contribution of predictors to the model to quantify the significance and applied the significant threshold as 1/number of predictors*100% ((*51*). Thus, the significant thresholds for BRT on species invasiveness (16 predictors) and community invasibility (12 predictors) were 6.26% and 8.33%, respectively.

As BRT accounts for spatial autocorrelation in neither the dependent nor predictor variables, we also ran an autoregressive error (SAR_error_) model for the community invasibility patterns and compared these results with those of the BRT, to check if spatial autocorrelation affected the results. We scaled all predictor variables to have zero mean and unit variance to ensure equal weighting in the model. Quadratic terms were included in the SAR_error_ model to consider non-linear responses. The spatial autocorrelation analysis was performed using the ‘spatialreg’ and ‘spdep’ packages (*52*). We used Nagelkerke’s R^2^ (*53*) as the pseudo R-squared to qualify the SAR_error_ model’s performance. After model fitting, we checked for broad spatial autocorrelation in model residuals by computing the Moran’s I statistic (*54*). The results of SAR_error_ models are provided in the supplementary material (Table S3). The core drivers identified by the BRT models were confirmed by the SAR_error_ analysis, suggesting spatial autocorrelation did not have an important effect on our results.

All statistical analyses were performed with R software version 4.1 (*55*).

## Author contributions

G.S., J.M.C. and S.B. designed the study, analyzed the data and wrote the manuscript. G.S. and A.M. collected and compiled the data, G.S. worked on the core data preparation and coding in R. All authors led to revising the paper and approving it for publication.

## Data and materials availability

All data needed to evaluate the conclusions in the paper are present in the paper and/or the Supplementary Materials. Additional data, scripts and files related to this paper will be available online after publication.

