## supplementary for "Species invasiveness and community invasibility of US freshwater fish fauna revealed via trait-based analysis"

**This PDF file includes:**

Figs. S1 to S5

Tables S1 to S3

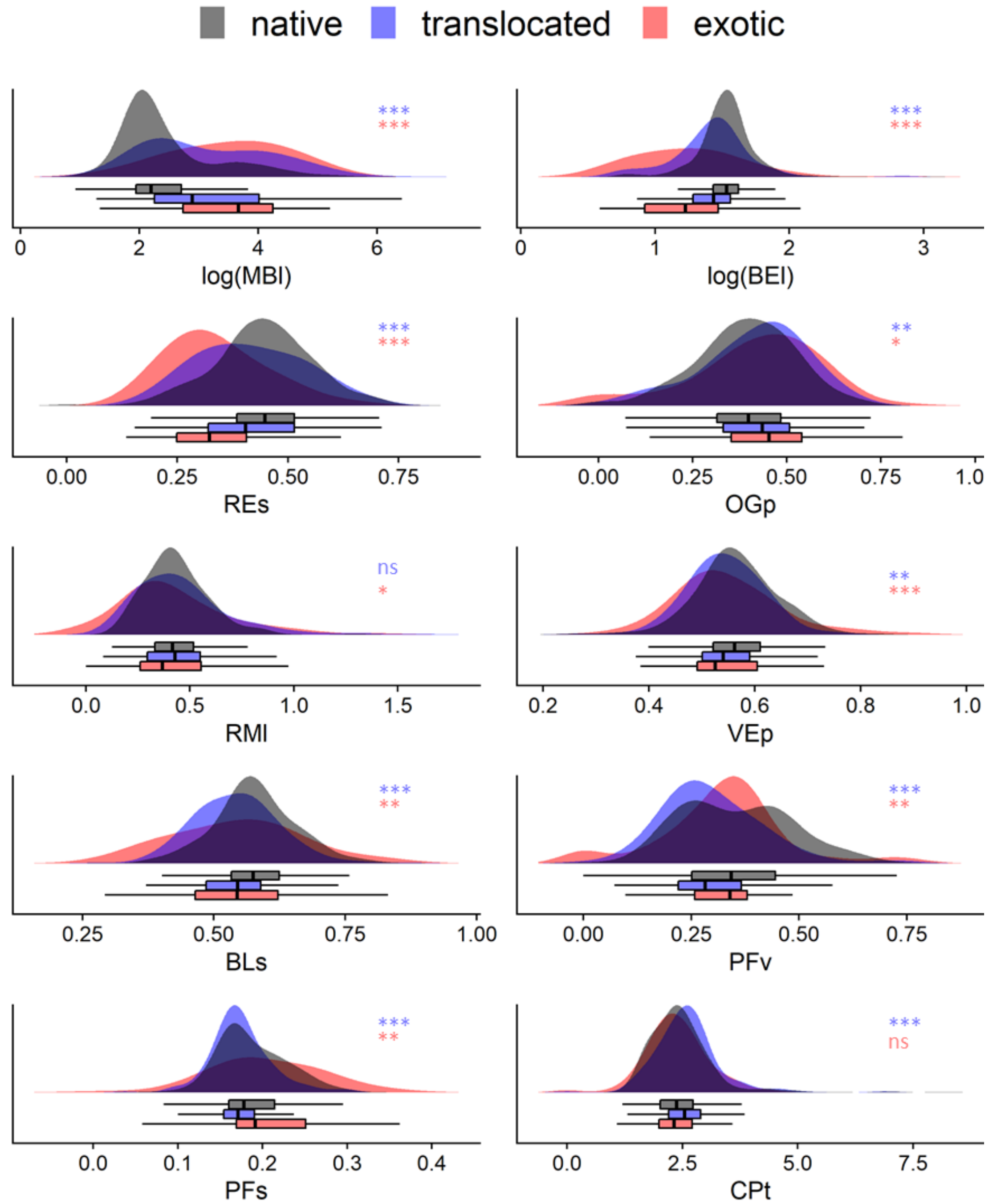


**Fig. S1.**

Density distribution of species on the 10 morphological trait axes for native, translocated and exotic species in the US. Boxplots and results of K-S tests of translocated-native (blue) and exotic-native (red) species are shown beside each plot item (*** *P* < .001, ** *P* < .01, * *P* < .05). Species number: native 562; translocated 227; exotic 73. See Table S2 for the complete results of K-S tests between the three groups. MBl: maximum body length; BEl: body elongation; REs: relative eye size; OGp: oral gape position; RMl: relative maxillary length; EVp: eye vertical position; BLs: body lateral shape; PFv: pectoral fin vertical position; PFs: Pectoral Fin Size; CPt: caudal peduncle throttling.

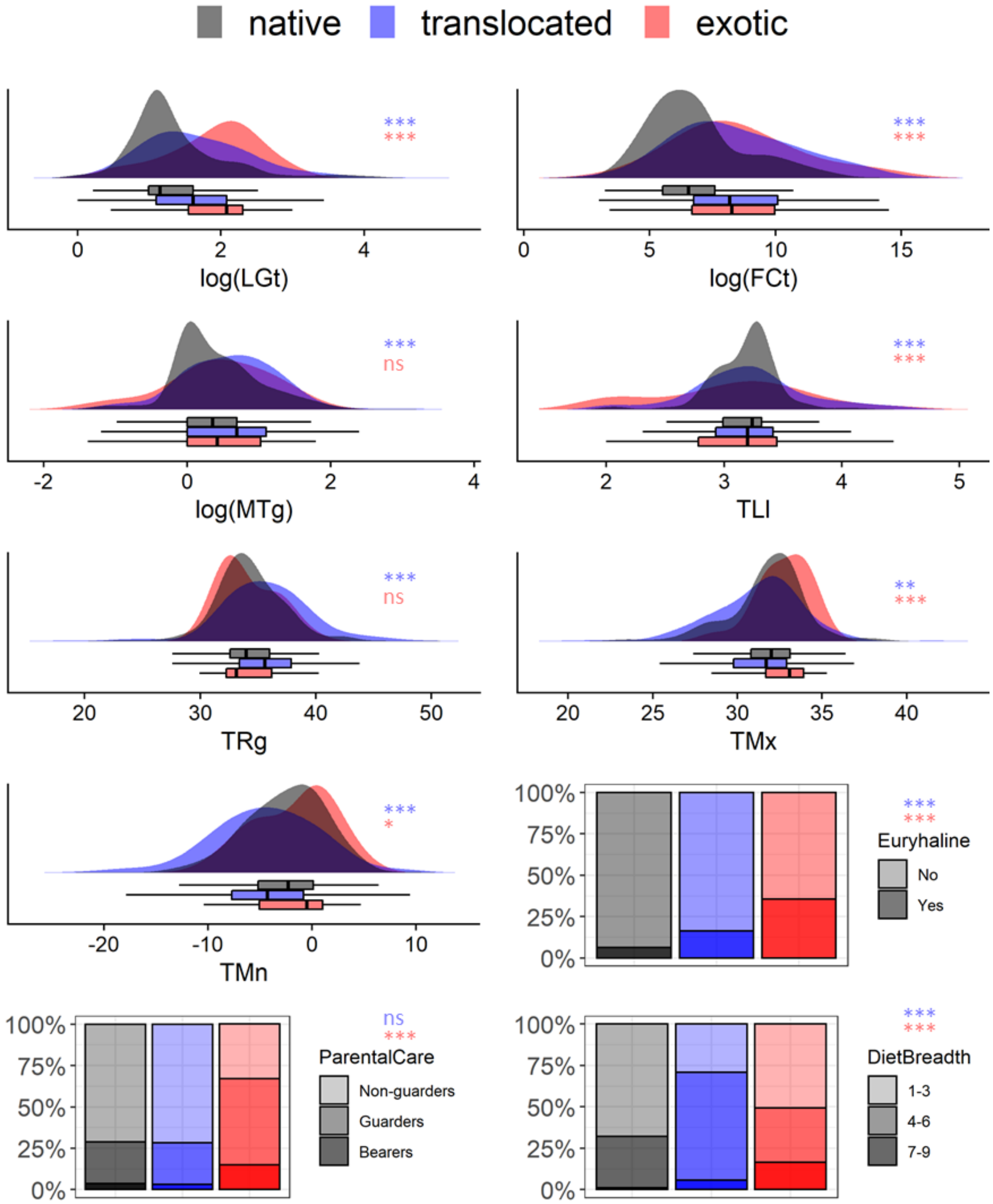


**Fig. S2.**

Density distribution of species on the other 10 life-historical trait axes for native, translocated and exotic species in US. Boxplots and results of K-S tests for the seven quantitative traits and Chi-square tests for the three categorical traits of translocated-native (blue) and exotic-native (red) species are shown beside each plot item (*** $P < .001$, ** $P < .01$, * $P < .05$). Species number: native 562; translocated 227; exotic 73. See Table S2 for the complete results of K-S tests for the first 7 quantitative traits and Chi-Square tests for the last 3 categorical traits between the three groups. LGt: longevity, FCt: fecundity; MTg: mature age; TLl: trophic level; TRg temperature range; TMx; maximum temperature; TMn: minimum temperature.

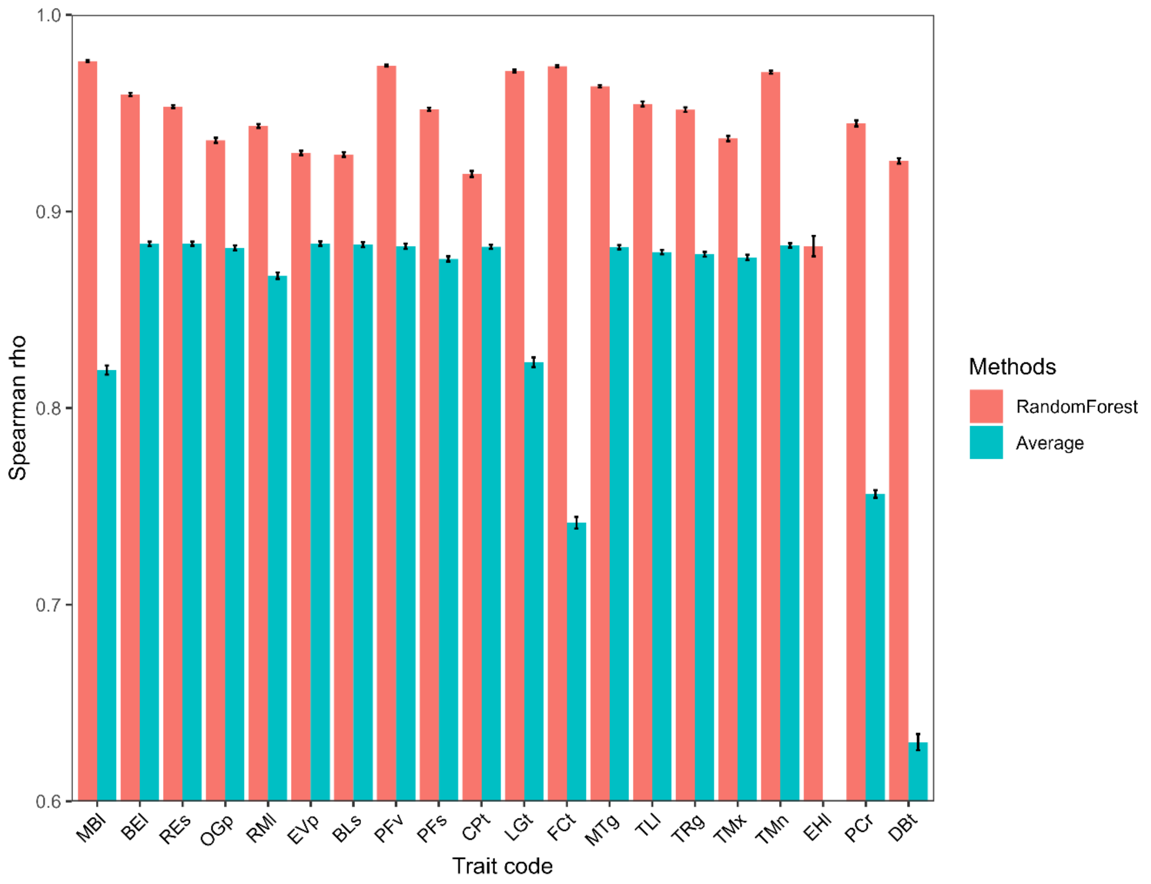


**Fig. S3.**

Comparison of the efficiency of filling the trait missing values using the random forest method and mean values. MBl: maximum body length; BEl: body elongation; REs: relative eye size; OGp: oral gape position; RMl: relative maxillary length; EVp: eye vertical position; BLs: body lateral shape; PFv: pectoral fin vertical position; PFs: Pectoral Fin Size; CPt: caudal peduncle throttling; LGt: longevity, FCt: fecundity; MTg: mature age; TLl: trophic level; TRg temperature range; TMx; maximum temperature; TMn: minimum temperature; EHl: euryhaline; PCr: parental care; DBt: diet breadth.

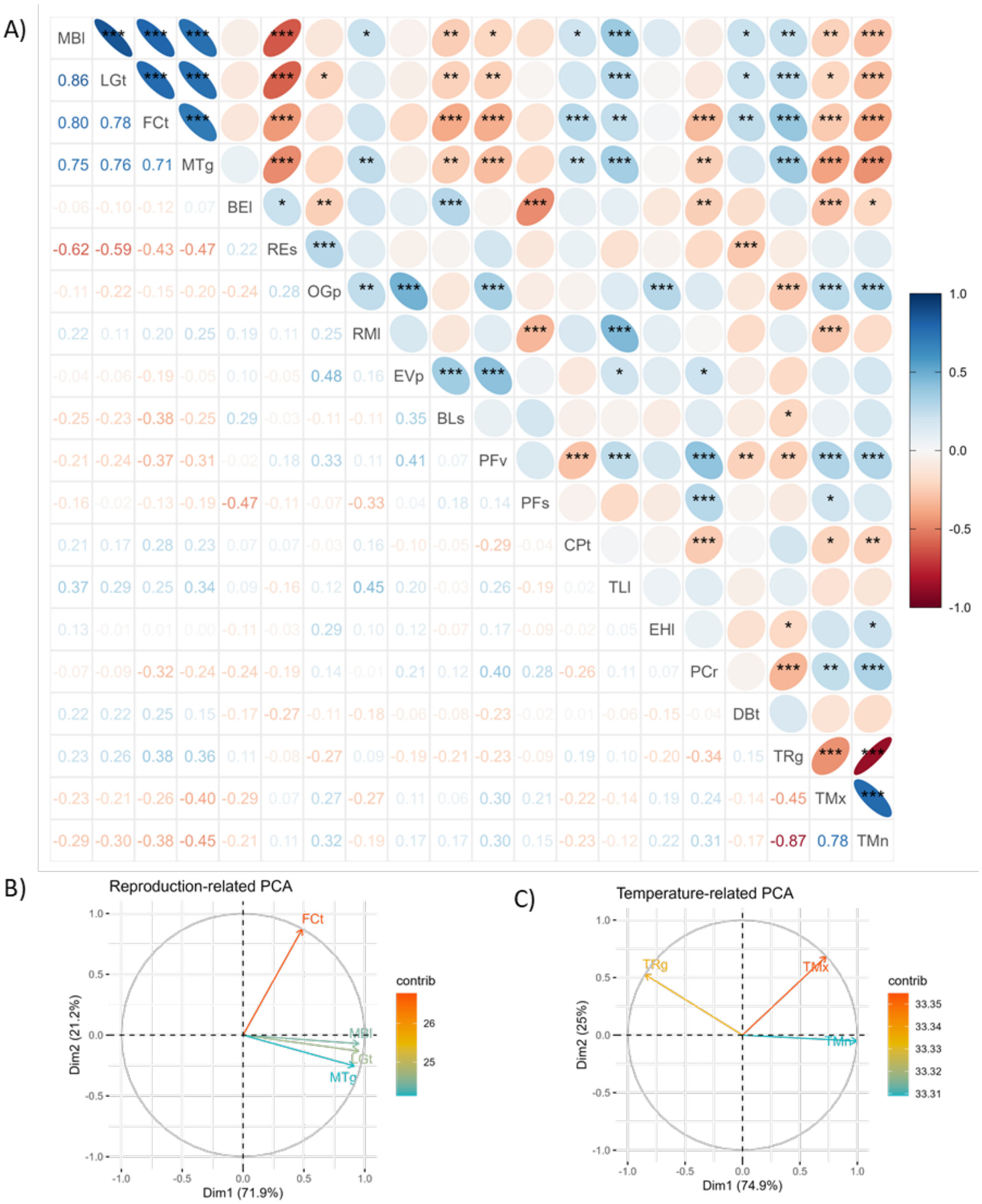


**Fig. S4.**

Relationship between the 20 functional traits for the 862 fish species in the US. A) Spearman correlation between the traits. B) PCA for the four reproduction-related traits. C) PCA for the three temperature-related traits. MBl: maximum body length; BEl: body elongation; REs: relative eye size; OGp: oral gape position; RMl: relative maxillary length; EVp: eye vertical position; BLs: body lateral shape; PFv: pectoral fin vertical position; PFs: Pectoral Fin Size; CPt: caudal peduncle throttling; LGt: longevity, FCt: fecundity; MTg: mature age; TLl: trophic level; TRg temperature range; TMx; maximum temperature; TMn: minimum temperature; EHl: euryhaline; PCr: parental care; DBt: diet breadth. (*** $P < 0.001$, ** $P < 0.01$, * $P < 0.05$)

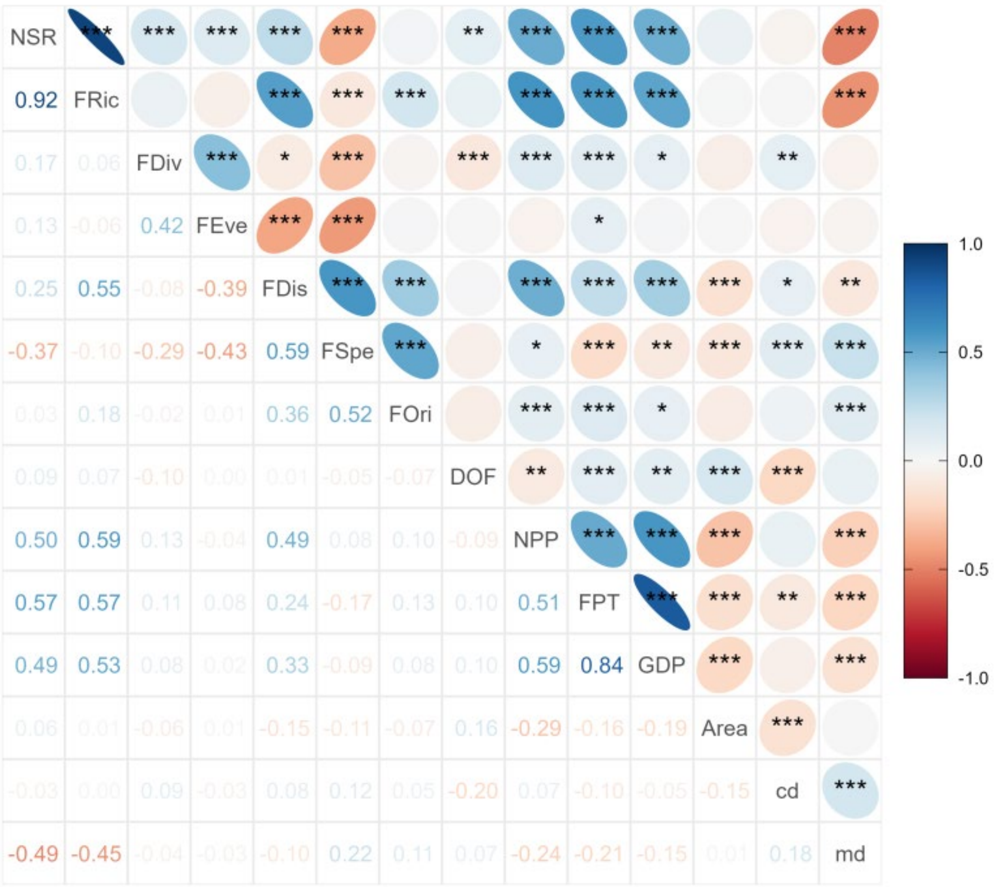


**Fig. S5.**

Spearman correlation between the 14 predictors considered to evaluate community invasibility (n = 1,873). NSR: native species richness; FRic: functional richness; FDiv: functional divergence; FEve: functional evenness; FDis: functional dispersion; FSpe: functional specialization; FOri: functional originality; DOF: degree of river fragmentation; NPP: net primary productivity; FPT: human footprint; GDP: gross domestic product; Area: watershed area; cd: centroid distance; md: mean distance. (*** $P < 0.001$, ** $P < 0.01$, * $P < 0.05$)

**Table S1.**

Type and source of the 20 functional traits of fish species in the United States.

| Trait Name | Abb. | Data type | Data source |
|---|---|---|---|
| **Morphological** | | | |
| Maximum Body Length | MBl | quantitative | FishBase |
| Body Elongation | BEl | quantitative | Su et al., 2021 |
| Relative Eye Size | REs | quantitative | Su et al., 2021 |
| Oral Gape Position | OGp | quantitative | Su et al., 2021 |
| Relative Maxillary Length | RMl | quantitative | Su et al., 2021 |
| Eye Vertical Position | EVp | quantitative | Su et al., 2021 |
| Body Lateral Shape | BLs | quantitative | Su et al.,2021 |
| Pectoral Fin Vertical Position | PFv | quantitative | Su et al., 2021 |
| Pectoral Fin Size | PFs | quantitative | Su et al., 2021 |
| Caudal Peduncle Throttling | CPt | quantitative | Su et al., 2021 |
| **Life-historical** | | | |
| Longevity | LGt | quantitative | Frimpong & Angermeier, 2010 |
| Fecundity | FCt | quantitative | Frimpong & Angermeier, 2010 |
| Mature age | MTg | quantitative | Frimpong & Angermeier, 2010 |
| Trophic Level | TLl | quantitative | FishBase |
| Temperature Range | TRg | quantitative | Frimpong & Angermeier, 2010 |
| Maximum Temperature | TMx | quantitative | Frimpong & Angermeier, 2010 |
| Minimum Temperature | TMn | quantitative | Frimpong & Angermeier, 2010 |
| Euryhaline | EHl | binary | Frimpong & Angermeier, 2010 |
| Parental Care | PCr | categorical | Frimpong & Angermeier, 2010 |
| Diet Breadth | DBt | categorical | Frimpong & Angermeier, 2010 |

**Table S2.**

Results of Kolmogorov–Smirnov tests for the seventeen quantitative traits and Chi-square tests for the three categorical traits between the native, translocated and exotic species on the 20 functional traits. Bold P-values are significant ($P < 0.05$).

| | P-value | | |
|---|---|---|---|
| Trait Name | Native-translocated | Native-exotic | Translocated-exotic |
| MBl | **0.000** | **0.000** | **0.007** |
| BEl | **0.000** | **0.000** | **0.000** |
| REs | **0.000** | **0.000** | **0.000** |
| OGp | **0.006** | **0.015** | 0.722 |
| RMl | 0.205 | **0.015** | 0.073 |
| EVp | **0.002** | **0.001** | 0.512 |
| BLs | **0.000** | **0.001** | **0.040** |
| PFv | **0.000** | **0.002** | **0.002** |
| PFs | **0.000** | **0.004** | **0.000** |
| CPt | **0.001** | 0.792 | **0.007** |
| LGt | **0.000** | **0.000** | **0.000** |
| FCt | **0.000** | **0.000** | 0.811 |
| MTg | **0.000** | 0.098 | 0.111 |
| TLl | **0.001** | **0.001** | 0.107 |
| TRg | **0.000** | 0.063 | **0.000** |
| TMx | **0.008** | **0.000** | **0.000** |
| TMn | **0.000** | **0.032** | **0.000** |
| EHl | **0.000** | **0.000** | **0.001** |
| PCr | 0.568 | **0.000** | **0.000** |
| DBt | **0.000** | **0.000** | **0.000** |

**Table S3.**

Results of spatially explicit simultaneous autoregressive (SAR) error models for the community invasibility to the established non-native species. Columns show the coefficient estimates (coef), standard error (SE), z-statistic value (z-value) and P-value for each variable. The significant P-values (<0.05) are in bold. The last two rows show the pseudo R-squared values and Moran's I statistic of the SAR model. ns: non-significant.

| Variable | coef | SE | z-value | *P*-value |
|---|---|---|---|---|
| FRic | 0.378 | 0.173 | 2.177 | **0.030** |
| FDiv | -0.676 | 0.138 | -4.896 | **0.000** |
| FEve | 0.470 | 0.146 | 3.219 | **0.001** |
| FDis | 0.318 | 0.252 | 1.259 | 0.208 |
| FSpe | 2.990 | 0.993 | 3.010 | **0.003** |
| FOri | -0.829 | 0.168 | -4.935 | **0.000** |
| DOF | 0.586 | 0.097 | 6.032 | **0.000** |
| NPP | -0.736 | 0.672 | -1.094 | 0.274 |
| FPT | 1.301 | 0.145 | 8.949 | **0.000** |
| Area | 2.154 | 0.157 | 13.727 | **0.000** |
| cd | -2.377 | 0.376 | -6.330 | **0.000** |
| md | 2.158 | 0.412 | 5.235 | **0.000** |
| $cd^2$ | 1.031 | 0.389 | 2.649 | **0.008** |
| $md^2$ | -1.187 | 0.418 | -2.836 | **0.005** |
| $NPP^2$ | 0.666 | 0.631 | 1.056 | 0.291 |
| $FSpe^2$ | -2.024 | 0.923 | -2.192 | **0.028** |
| $Area^2$ | -1.152 | 0.142 | -8.143 | **0.000** |
| pseudo R-squared | | 0.508 | | |
| Moran's I statistic | | -0.0431 (ns) | | |
